# H3 K27M and EZHIP impede H3K27-methylation spreading by inhibiting allosterically stimulated PRC2

**DOI:** 10.1101/2020.08.05.236737

**Authors:** Siddhant U. Jain, Andrew Q. Rashoff, Samuel D. Krabbenhoft, Dominik Hoelper, Truman J. Do, Tyler J. Gibson, Stefan M Lundgren, Eliana R. Bondra, Shriya Deshmukh, Ashot S. Harutyunyan, Nikoleta Juretic, Nada Jabado, Melissa M. Harrison, Peter W. Lewis

**Affiliations:** Department of Biomolecular Chemistry, School of Medicine and Public Health, University of Wisconsin, Madison, WI 53715, USA; Department of experimental Medicine, McGill University, Montreal, QC, Canada; Department of Human Genetics, McGill University, Montreal, QC H3A 1B1, Canada. Department of Pediatrics, McGill University, and The Research Institute of the McGill University Health Center, Montreal, QC H4A 3J1, Canada

## Abstract

Diffuse midline gliomas and posterior fossa type-A ependymomas contain the highly recurrent histone H3 K27M mutation and the H3 K27M-mimic EZHIP, respectively. *In vitro*, H3 K27M and EZHIP are competitive inhibitors of Polycomb Repressive Complex 2 (PRC2) lysine methyltransferase activity. *In vivo*, these proteins reduce overall H3K27me3 levels, however residual peaks of H3K27me3 remain at CpG islands through an unknown mechanism. Here, we report that EZHIP and H3 K27M preferentially interact with an allosterically activated form of PRC2 *in vivo*. The formation of H3 K27M- and EZHIP-PRC2 complexes occurs at CpG islands containing H3K27me3 and impedes PRC2 and H3K27me3 spreading. While EZHIP is not found outside of placental mammals, we find that expression of human EZHIP reduces H3K27me3 in *Drosophila melanogaster* through a conserved molecular mechanism. Our results highlight the mechanistic similarities between EZHIP and H3 K27M *in vivo* and provide mechanistic insight for the retention of residual H3K27me3 in tumors driven by these oncogenes.

---

Diffuse midline gliomas (DMGs) are highly aggressive pediatric tumors with a poor prognosis. About ~80% of DMGs contain the recurrent histone H3 lysine 27 (H3 K27M) mutation ^1,2^. Despite representing a small fraction of total histone H3 (3-17%), H3 K27M causes a global reduction of H3 lysine 27 trimethylation (H3K27me3) in DMGs ^3–8^. The K27M mutation occurs in genes encoding histone H3 proteins that are assembled into nucleosomes through replication-coupled (H3.1/2) and replication-independent mechanisms (H3.3). Posterior fossa type-A ependymomas (PFA ependymomas) display similar gene expression, DNA methylation, and low H3K27me3 profiles as DMGs ^9^. However, instead of recurrent H3 K27M mutations, PFA ependymomas aberrantly express a newly discovered gene, *EZHIP* (also known as *CXorf67, CATACOMB*) ^10^. *EZHIP-expressing* ependymomas are generally found in young children (median age 3 years) and display poorer prognosis as compared to other posterior fossa ependymomas ^11^. While numerous studies have described the biochemical activities of H3 K27M oncohistones, their precise role and function in modulating gene expression remains unclear. Moreover, the molecular mechanisms of EZHIP in cells remains largely uncharacterized.

Polycomb Repressive Complex 2 (PRC2) catalyzes H3K27me3, which is involved in transcriptional silencing. The oncoproteins EZHIP and H3 K27M are competitive inhibitors of PRC2 ^12–14^. A C-terminal peptide within EZHIP mimics the H3 K27M sequence and is necessary and sufficient to inhibit PRC2 activity *in vitro*. Expression of EZHIP or H3 K27M in cells leads to global reduction of H3K27me2/3 ^12–15^. However, hundreds of CpG islands (CGIs), that represent high-affinity PRC2-binding sites retain residual H3K27me3 ^12,16–18^. Residual PRC2 activity is necessary for survival of H3 K27M-containing gliomas ^16–18^. It has been proposed that retention of H3K27me3 at CGIs is necessary to silence tumor suppressor genes and maintain cell proliferation. Nonetheless, the molecular mechanism by which EZHIP and H3 K27M reduce H3K27me3 specifically from intergenic regions but disproportionately retain H3K27me3 at CGIs remains elusive.

PRC2 interacts with unmethylated CGIs through its auxiliary subunits, Polycomb-like proteins (PCLs) or JARID2, where it catalyzes high levels of H3K27me3 ^19–25^. H3K27me3, initially catalyzed at CGIs, interacts with the EED subunit and allosterically stimulates PRC2 catalytic activity by ~8-fold ^26,27^. In addition to H3K27me3, trimethylation of K116 in the JARID2 subunit can also stimulate EZH2 activity in an EED-dependent manner ^28^. This ‘read-write’ mechanism triggers PRC2 spreading into the intergenic regions and formation of broad H3K27me3 domains. Remarkably, the inhibitory potential of H3 K27M and EZHIP oncoproteins is substantially enhanced by allosteric stimulation of PRC2 *in vitro* ^12,29–31^. It is unclear whether the preferential inhibition of allosterically stimulated PRC2 by EZHIP and H3 K27M plays a functional role *in vivo*.

Here, we demonstrate that EZHIP preferentially interacts with allosterically stimulated PRC2 *in vitro* and *in vivo*. EZHIP impedes PRC2 spreading by stabilizing a high-affinity complex between H3K27me3-PRC2-EZHIP at CGIs containing residual H3K27me3. Using reChIP experiments, we demonstrate that H3 K27M oncohistones interact with and stall PRC2 spreading at CGIs. Finally, despite its absence in non-placental mammals, we demonstrate that EZHIP reduces H3K27me3 in *Drosophila melanogaster* through a conserved molecular mechanism. In summary, we provide evidence that EZHIP and H3 K27M bind PRC2 *in vivo* and reduce H3K27me2/3 in trans by blocking PRC2 spreading.

## Results

### EZHIP preferentially interacts with allosterically stimulated PRC2 *in vivo*

Expression of *EZHIP* in cells leads to an overall reduction of H3K27me3, however residual H3K27me3 is retained at CGIs (Jain et al., 2019). The mechanism by which these narrow H3K27me3 peaks persists in tumors expressing *EZHIP* remains elusive. Therefore, we used ChIP-Sequencing (ChIP-seq) to identify the genomic regions where EZHIP inhibits PRC2 in mouse embryonic fibroblasts expressing EZHIP. Surprisingly, we found that EZHIP and EZH2 colocalized with residual H3K27me3 at CGIs (Figure 1A-D). Depletion of EED through transient expression of Cre-recombinase abolished EZHIP enrichment suggesting that EZHIP binds to chromatin indirectly through its interaction with PRC2 (Figure 1A, B; S1A, B). A K27M-like peptide (KLP) within the C-terminus of EZHIP interacts with EZH2 catalytic site residues (where M406 is equivalent to H3 K27M) ^12^. Consequently, EZHIP M406E mutant within the KLP failed to inhibit PRC2 *in vitro* and did not immunoprecipitate PRC2 subunits from nuclear extract (Figure 1E, F; S1A). Moreover, we did not identify enrichment of EZHIP M406E at CGIs with PRC2 and H3K27me3 *in vivo* (Figure 1G; S1C). These data linked *in vitro* PRC2 inhibition and *in vivo* reduction of H3K27me3 to interaction between EZHIP KLP and EZH2 at CGIs.

**Figure 1.**
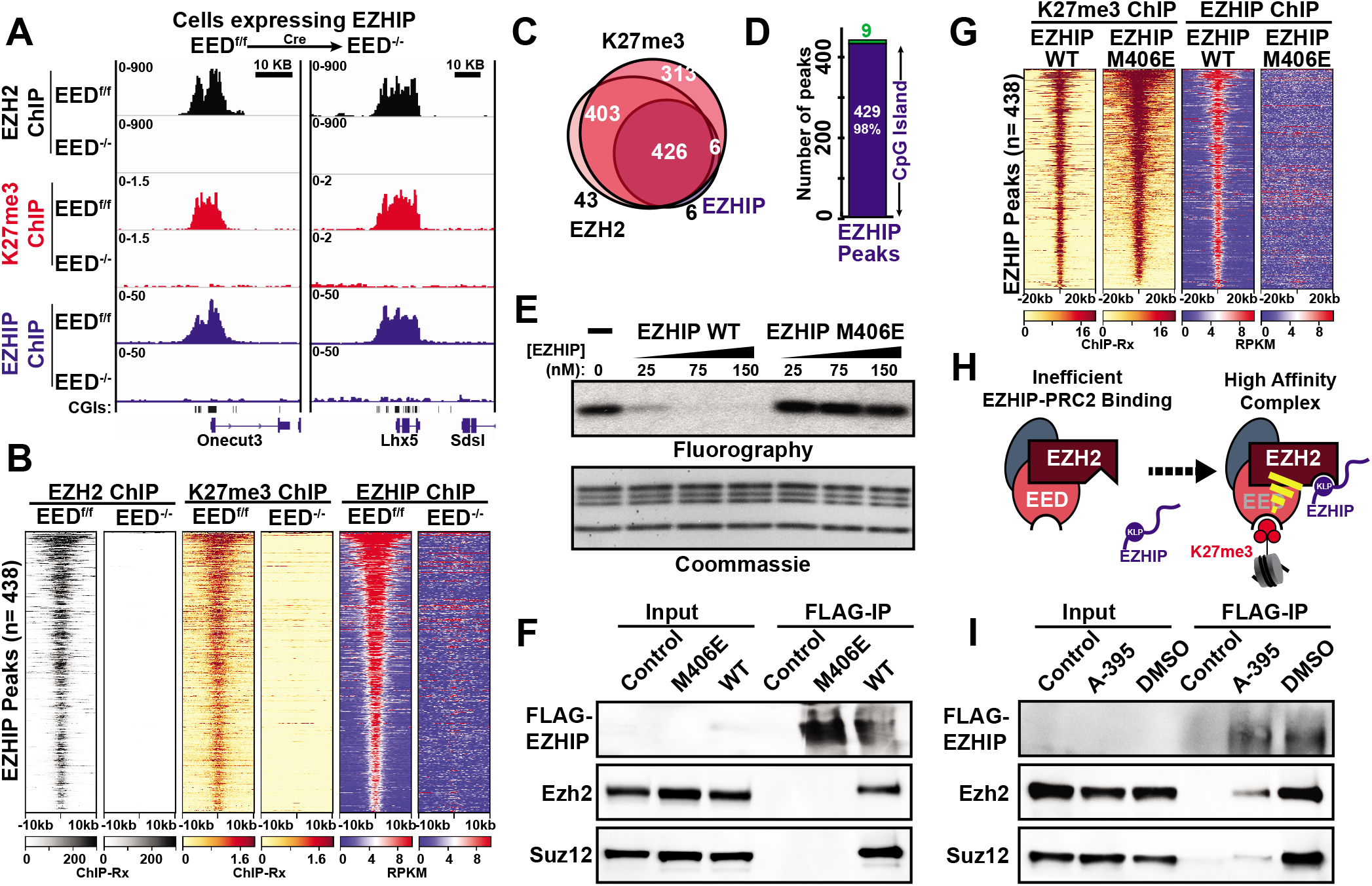
EZHIP preferentially intereacts with allosterically stimulated PRC2. **(A)** EZH2, H3K27me3 and EZHIP ChIP-Seq enrichments in EEF^f/f^ or EED^-/-^ MEFs expressing FLAG-tagged EZHIP. **(B)** EZH2, H3K27me3 and EZHIP ChIP enrichments in EEF^f/f^ or EED^-/-^ MEFs expressing *EZHIP* at EZHIP peaks. **(C)** Overlap between EZHIP, H3K27me3 and EZH2 peaks in MEFs expressing *EZHIP*. **(D)** Fraction of EZHIP peaks that contain CpG islands (blue). **(E)** HMT assays: 20 nM PRC2 was incubated with 200 nM nucleosomes, 4 μM S-adenosyl Methionine and 1 μM ^3^H-S-adenosyl Methionine. EZHIP WT or M406E was titrated into the reaction mixture. PRC2 activity on histone H3 was detected using fluorography following SDS-PAGE. **(F)** Immunoblots of inputs and elutions from FLAG-affinity purification of EZHIP WT or M406E from MEFs. **(G)** Normalized enrichment of H3K27me3 and FLAG-EZHIP ChIPs from MEFs expressing EZHIP WT or M406E mutant. (H) Binding of H3K27me3 to EED leads to a conformational change in PRC2, which increases its affinity for EZHIP. **(I)** Immunoblots of inputs and elutions from FLAG-affinity purification of EZHIP WT from MEFs in the presence or absence of EED inhibitor, A-395.

Previously, we showed that the PRC2 inhibitory potential of EZHIP is significantly enhanced in the presence of H3K27me3 peptide ^12^. The interaction between EED and H3K27me3 is proposed to induce or stabilize an EZH2 conformation that has increased affinity towards the peptide substrates and inhibitors, such as EZHIP and H3 K27M (Figure 1H). We assessed EZHIP-PRC2 interactions using FLAG affinity purification in the presence or absence of a H3K27me3-competitive inhibitor of EED, A-395 ^32^. Cells treated with A-395 exhibit low H3K27me3 levels due to reduced allosteric activation of PRC2 ^32^. We found that EZHIP immunoprecipitated substantially lower amounts of PRC2 subunits from MEF nuclear extracts in the presence of A-395 (Figure 1I). These data suggest that EZHIP preferentially interacts with allosterically stimulated PRC2 and help explain why the inhibitory protein EZHIP interacts with PRC2 at regions containing H3K27me3 *in vivo*.

### EZHIP reduces PRC2 spreading by stalling it at CpG islands

We hypothesize that the formation of a catalytically inactive ternary complex, H3K27me3-PRC2-EZHIP, restrains PRC2 from spreading into the intergenic regions and hence, increases PRC2 residency at CGIs containing H3K27me3 (Figure 2A; S2A). Using quantitative ChIP-Seq, we found that expression of wildtype EZHIP, but not the PRC2-binding deficient M406E mutant, led to a substantial increase of EZH2 occupancy at residual H3K27me3 sites containing high CpG density (Figure 2B, C; S2B-E). Simultaneously, EZH2 enrichment was significantly reduced from CpG-poor domains of H3K27me3 (Figure S2C, D), suggesting that EZHIP stalls PRC2 at CGIs that serve as PRC2 recruitment sites. To test our model in human cancers, we used SUZ12 CUT&RUN data from U2OS osteosarcoma cells that express endogenous EZHIP ^15^. Mirroring our results in MEFs expressing *EZHIP*, wildtype U2OS cells exhibit sharp SUZ12 peaks, but these peaks redistributed to broader intergenic domains upon loss of *EZHIP* (Figure 2D, E; S2F, G). These results indicate that EZHIP impedes PRC2 spreading by stalling it at recruitment sites.

**Figure 2.**
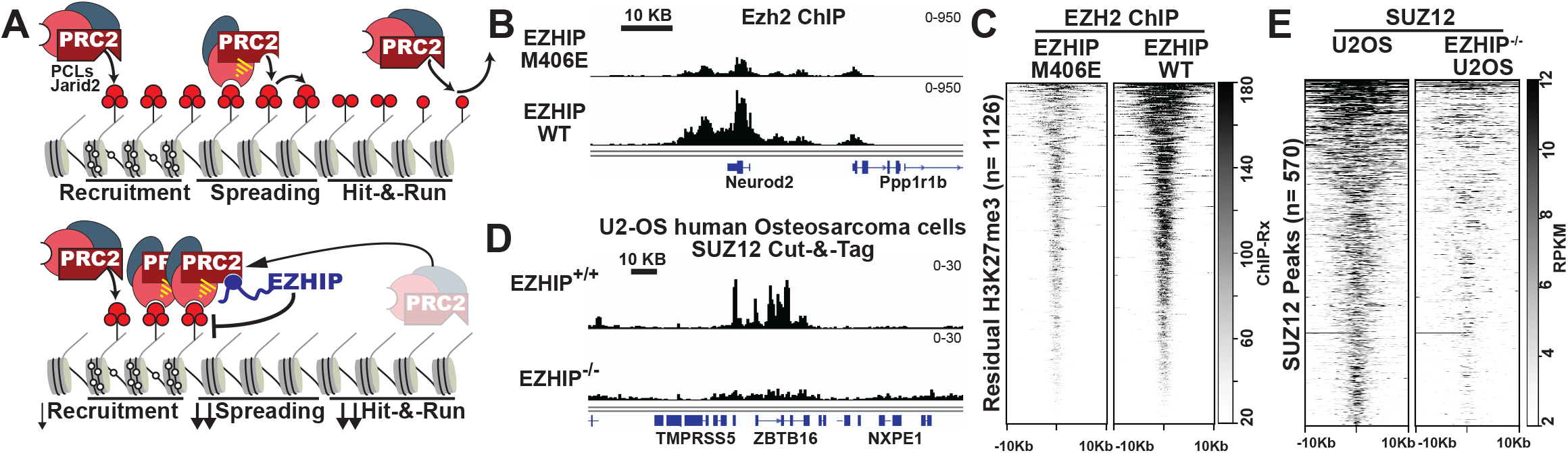
EZHIP inhibits PRC2 spreading by stalling it at CGIs containing residual H3K27me3. **(A)** EZHIP-mediated re-distribution of PRC2. Normally, PRC2 catalyzes H3K27 methylation on upto 90% of total Histone H3. Therefore, only a small fraction of PRC2 is present at recruitment sites (CGIs). In cells expressing *EZHIP*, EZHIP stalls PRC2 spreading by forming a high affinity complex with allosterically stimulated PRC2 at recruitment sites containing residual H3K27me3 and thereby, increasing its steady state occupancy at CGIs. (**B)** EZH2 ChIP-Rx profiles in MEFs expressing EZHIP WT or M406E. (**C**) EZH2 ChIP-Rx enrichment centered at residual H3K27me3 peaks. **(D)** SUZ12 CUT&RUN occupancy (RPKM) in U2OS cells expressing endogenous *EZHIP* or *EZHIP*^/-^ cells. **(E)** SUZ12 enrichment (RPKM) at SUZ12 peaks in U2OS cells.

### H3 K27M mutations interact with and stall PRC2 at CpG island

EZHIP inhibits PRC2 through a mechanism that is similar to H3 K27M oncohistones and leads to PRC2 redistribution from intergenic regions to CGIs by interacting with allosterically stimulated PRC2. Nevertheless, whether H3 K27M directly interacts with PRC2 *in vivo* at steady state has remained controversial because H3 K27M and PRC2 occupancies do not correlate positively in cells ^17,18,30^. The amount of H3 K27M far exceeds that of PRC2 in the nucleus ^30^. H3 K27M, being histones, are incorporated into nucleosomes and are therefore present throughout the genome. In contrast, PRC2 is localized in relatively narrow peaks at CGIs, and this may explain poor correlation between the enrichment of H3 K27M and PRC2. Previously, we and others have found that H3 K27M preferentially binds and inhibits PRC2 in the presence of H3K27me3, similar to EZHIP ^12,29–31^. Therefore, we hypothesize that H3 K27M also stalls PRC2 at CGIs containing residual H3K27me3 by binding allosterically stimulated PRC2.

To directly map genomic regions where H3 K27M interacts with PRC2 in cells, we used H3 K27M ChIP followed by EZH2 reChIP-Seq (Figure 3A; S3A, B). We selected reads with fragment size smaller than 400 bp for our analyses to only capture PRC2 bound to mono- or dinucleosomes. As would be predicted from our model, H3 K27M-bound EZH2 was enriched at CGIs containing residual H3K27me3 (Figure 3B-E; S3C, D). We did not detect EZH2 reChIP enrichment in control cells that did not express a FLAG-tagged H3 K27M transgene, confirming our detection of only H3 K27M-bound EZH2 instead of a background signal (Figure S3C). These data suggest that H3.1 and H3.3 K27M directly interact with PRC2 at CGIs, similar to EZHIP.

**Figure 3.**
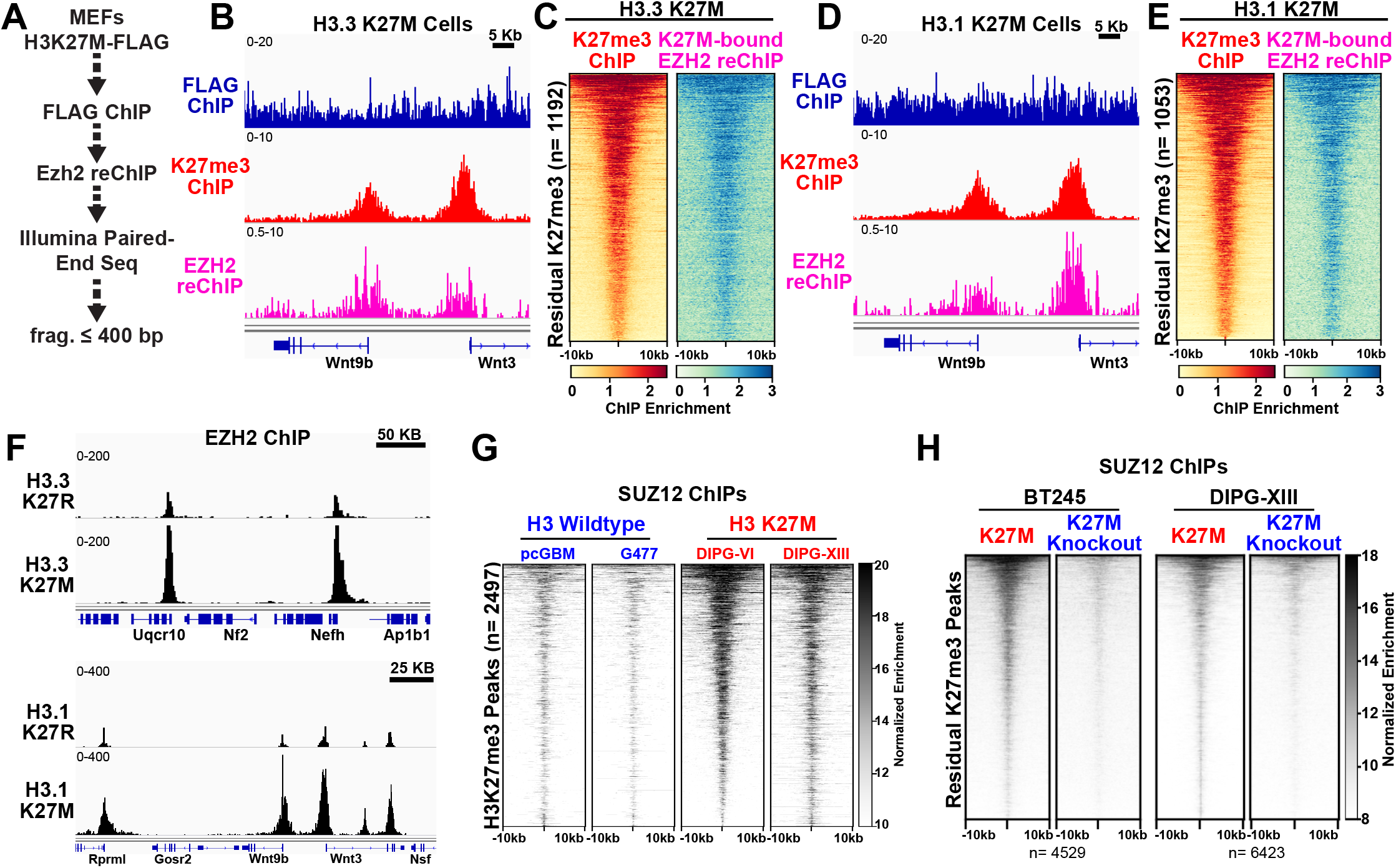
H3 K27M reduces H3K27me3 in *trans* by stalling PRC2 at CpG islands. **(A)** Sequential ChIP-Seq used to identify genomic localtions of H3K27M-bound Ezh2. **(B)** H3.3 K27M-FLAG ChIP, H3K27me3 ChIP and Ezh2 reChIP in cells expressing FLAG-tagged H3.3 K27M. **(C)** H3K27me3 and EZH2 reChIP profiles in cells expressing H3.3K27M at residual H3K27me3 peaks. **(D and E)** Same as **B** and **C** respectively but for cells expressing FLAG-tagged H3.1K27M. **(F)** Ezh2 ChIP-Rx enrichment in cells expressing H3.3 (top) or H3.1 (bottom) K27M or K27R. **(G)** SUZ12 ChIP enrichment in DIPGs containing H3K27M mutations or H3WT at combined H3K27me3 peaks (Harutyunyan et al 2019). **(H)** Heatmap displaying SUZ12 enrichment in DIPGs cell lines containing H3K27M mutation or corrosponding H3K27M knockout cells.

Next, we examined changes in PRC2 distribution in cells expressing H3 K27M by mapping overall EZH2 binding profile. Similar to cells expressing *EZHIP*, expression of H3.1 or H3.3 K27M led to a significant increase in EZH2 occupancy at CGIs containing residual H3K27me3 and a concurrent reduction of EZH2 from non-CpG-rich regions (Figure 3F; S3F-K). To corroborate our model in gliomas, we profiled genomic distribution of SUZ12 in patient-derived DMG cell lines with wildtype H3 or H3 K27M mutations ^18^. DMGs lines containing the H3 K27M mutation had significantly higher SUZ12 enrichment at CGIs relative to H3 wildtype glioma lines (Figure 3G; S4A-D). Importantly, Cas9-mediated genetic ablation of H3.3 K27M substantially reduced SUZ12 occupancy at these sites (Figure 3H, S4E-K). Taken together, our results suggest that H3 K27M directly interacts with PRC2 *in vivo* and stalls it at CGIs containing residual H3K27me3.

### Mammalian EZHIP inhibits Drosophila PRC2 through a conserved mechanism

Our studies demonstrate remarkable similarities in the mechanism through which EZHIP and H3 K27M inhibit PRC2 through directly interacting with it at its recruitment sites. While histone H3 is highly conserved among eukaryotes, EZHIP is only present in placental mammals. Previous studies have found that expression of H3 K27M in fruit flies largely phenocopies loss of PRC2 activity ^33^. Since EZHIP mimics the molecular function of H3 K27M oncohistone, we hypothesized that human EZHIP may inhibit *Drosophila* PRC2 despite its evolutionary absence in flies. Indeed, we found that expression of human EZHIP or H3.3 K27M in imaginal wing discs led to substantial reduction of H3K27me3, relative to EZHIP M406E or H3 K27R controls (Figure 4A). These data further highlight the similarity between the molecular functions of the two oncogenes.

**Figure 4.**
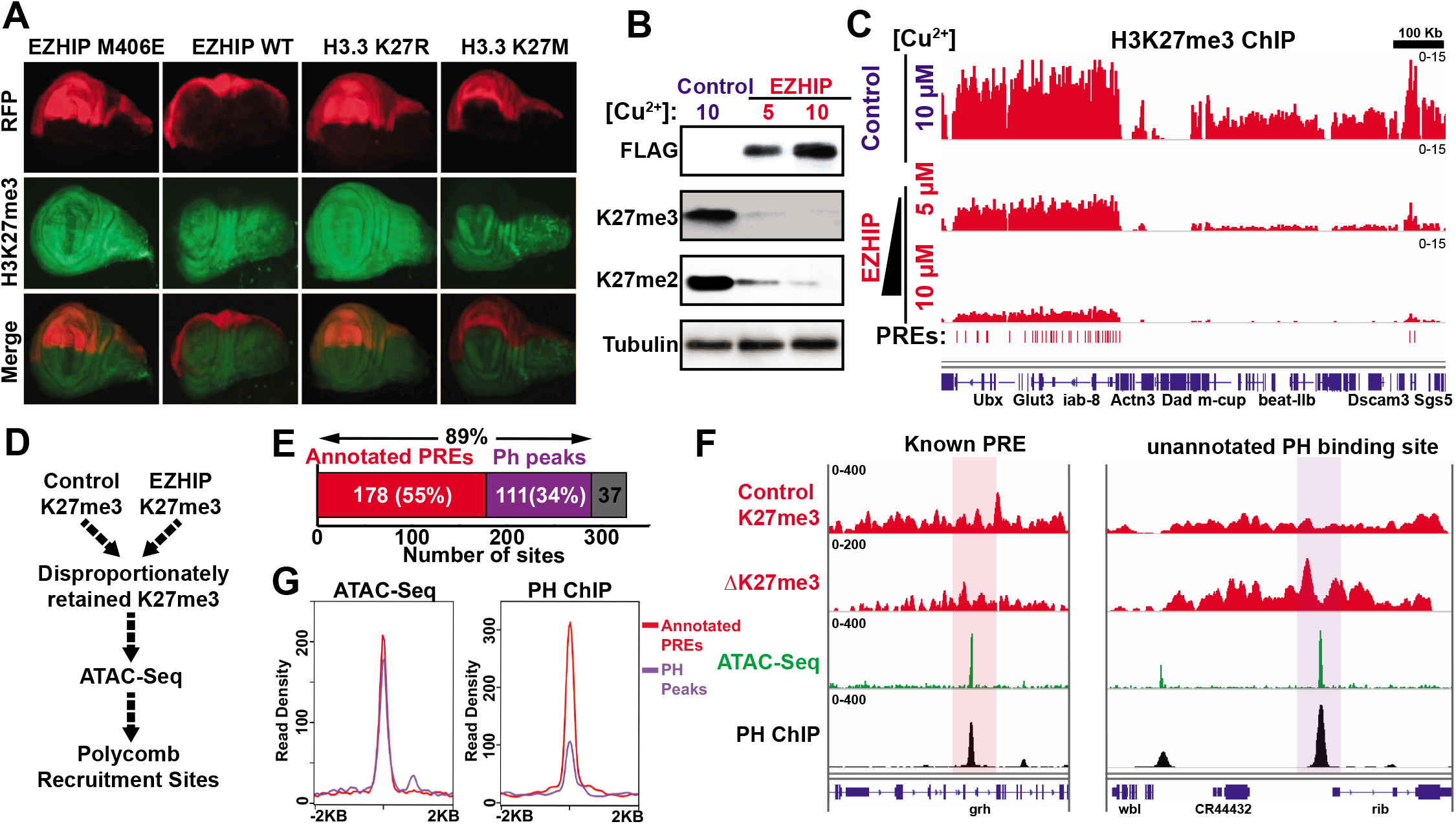
Human EZHIP reduces H3K27me3 in Drosophila through a conserved mechanism. **(A)** H3K27me3 staining (green) of wing imaginal discs from third instar larvae expressing either EZHIP WT, EZHIP M406E, H3 K27M, or H3 K27R (as indicated) driven by *en-GAL4*. RFP (red) indicates the region of *en-GAL4* expression. **(B)** Immunoblots of whole cell extracts from S2 cells expressing FLAG-tagged EZHIP induced with 5 μM or 10 μM copper for 72 hours. Empty vector was used as control. **(C)** Reference-normalized H3K27me3 profile in S2 cells as described in **B**. **(D)** Strategy to identify putative PRC2-recruitment sites within residual H3K27me3 peaks in S2 cells. Genomic regions that disproportionately retained H3K27me3 in cells expressing EZHIP were determined as a difference of internally-normalized H3K27me3 (EZHIP-Control). **(E)** Fraction of putative PRC2 recruitment sites within residual H3K27me3 that contained annotated Polycomb Response Elements (PREs) or Ph peaks. **(F)** Baseline H3K27me3 ChIP, ΔH3K27me3 (EZHIP-Control), ATAC-Seq and PH ChIP-Seq profiles (related to figure **S7C**). Previously annotated PREs and other PRC2 recruitment sites are highlighted in red and purple respectively. **(G)** ATAC-Seq and Ph ChIP-Seq densities at putative PRC2 recruitment sites.

The *cis*-regulatory elements involved in PRC2 recruitment that account for the global H3K27 methylation profile have not been identified in mammals. However, PRC2 is recruited to Polycomb Response Elements (PREs) through its Pho subunit in *Drosophila melanogaster*^34,35^. Therefore, *Drosophila* presents an excellent model to study and validate PRC2 recruitment versus spreading defects mediated by EZHIP. Having validated the ability of EZHIP to inhibit H3K27me3 *in vivo*, we established a copper-inducible system to express EZHIP in *Drosophila* S2 cells and showed that expression of EZHIP led to a dose-dependent reduction of H3K27me3, consistent with a competitive mode of inhibition with respect to histone substrates (Figure 4B, C). Similar to EZHIP in mammalian cells, ChIP-seq for H3K27me3 identified regions that retained residual levels of H3K27me3 upon EZHIP induction in S2 cells (Figure 4C; S5A, B). Furthermore, majority of the PREs retained residual H3K27me3 in cells expressing EZHIP (Figure 4C, S5B).

To systematically distinguish between PRC2 recruitment versus spreading defects, we identified genomic regions that disproportionately retained H3K27me3 relative to global H3K27me3 (Figure 4D). Because PREs are characterized by being nucleosome-free regions, we used ATAC-Seq to identify potential PRC2 recruitment sites within the residual H3K27me3 regions. We found that 178/326 (55%) of the regions that retained H3K27me3 contained previously annotated PREs and displayed polyhomeotic (PH) enrichment (Figure 4E-G, S5C, D). Moreover, an additional 111 (34%) sites also displayed PH occupancy, accounting for ~90% of regions that retained H3K27me3 in cells expressing EZHIP (Figure 4E-F). Notably, the amplitude of PH enrichment at unannotated PH binding sites was lower as compared to annotated PREs (Figure 4G), likely representing cell-type specific, weaker PRC2-recruitment sites ^36^. These results further support the model that EZHIP preferentially inhibits PRC2 spreading. Moreover, this model provides a mechanistic explanation for the presence of sharp H3K27me3 peaks in DMGs and PFA ependymomas at genes containing CGIs including the tumor suppressor, *CDKN2A*.

## Discussion

In this study, we used genomic approaches to demonstrate that glioma-driving oncoproteins EZHIP and H3 K27M directly interact with PRC2 *in vivo* and reduce H3K27me3 in *trans*. Due to the broad chromatin deposition profile of histone H3, it had been challenging to assess PRC2-H3 K27M interaction *in vivo* using such approaches. We show that the non-histone, H3 K27M-mimic EZHIP occupies the same sites as PRC2 *in vivo*. Importantly, we successfully detected H3 K27M-PRC2 interaction *in vivo* using a sequential ChIP strategy. These data support the model that H3 K27M inhibits PRC2 activity in *trans* and link the numerous studies that characterized the PRC2-K27M interactions *in vitro* to the *in vivo* loss of H3K27me3.

Previous studies revealed that H3 K27M and EZHIP have substantially higher affinity for allosterically stimulated PRC2. In the complete absence of H3K27me3, EZHIP and H3 K27M appear to be poor inhibitors of PRC2 and hence, allow initial H3K27me3 at high-affinity recruitment sites ^37,38^. However, these oncoproteins form high-affinity complexes with H3K27me3-bound PRC2 at these recruitment sites to prevent further catalysis of H3K27me3 at distal sites. As a result, PRC2-recruitment sites disproportionately retain H3K27me3 at equilibrium in these tumors. We propose that tumors expressing EZHIP or H3 K27M retain residual H3K27me3 at PRC2 recruitment sites because of their preferential interaction with allosterically stimulated PRC2.

We found that EZHIP and H3 K27M bind to allosterically stimulated PRC2 at CGIs containing residual H3K27me3. While we focused on H3K27me3-stimulated PRC2 in this study, it is known that Jarid2 K116me3 also stimulates PRC2 by interacting with EED. Additionally, Jarid2 serves to target PRC2 at its recruitment sites through an independent mechanism ^19,20,22,25,28,39^. Therefore, enhanced binding of EZHIP and H3 K27M to PRC2 stimulated by Jarid2-K116me3 may also contribute to increased PRC2 residence at CGIs and stalling its spread.

Recurrent H3 K27M mutations and aberrant *EZHIP* expression are frequently found in DMGs and PFA ependymomas, respectively. Our study suggests that EZHIP and H3 K27M inhibit PRC2 activity in *trans*, through a similar molecular mechanism *in vitro* and in *vivo*. Remarkably, two recent studies discovered aberrant expression of *EZHIP* in a subset of DMGs lacking H3 K27M mutations ^40,41^. Similarly, a small fraction of PFA ependymomas contain H3 K27M mutations that are mutually exclusively with *EZHIP* expression ^10^. Our finding that EZHIP and H3 K27M have similar underlying biochemical mechanisms are consistent with the clinical observations that both oncoproteins appear to drive the same subtype of gliomas. Moreover, pharmacological interventions that have been proposed for H3 K27M-positive gliomas might be promising candidates in gliomas expressing *EZHIP*^8,16,17,42–44^.

We validated our findings that EZHIP disproportionately blocks PRC2 spreading while sparing residual H3K27me3 at recruitment sites using *Drosophila melanogaster* S2 cells. Half of the sites that disproportionately retained H3K27me3 were previously characterized *cis*-acting PREs. However, we also found more than one hundred new, weak PRC2-binding sites that likely represent tissue-specific PREs. Previous studies used a combination of H3K27me3 ChIP- and ATAC-Seq to identify PREs in *Drosophila melanogaster*. Expression of EZHIP might provide a tool to filter out majority of the genomic regions containing H3K27me3 and a more sensitive method to detect tissue specific PREs in future studies.

## Methods

### Transgenic cell lines and culture

Mouse embryonic fibroblasts used in this study containing loxP sites flanking exon 3-6 of EED were described previously ^12^. Cells were cultured in DMEM supplemented with 10% FBS, 1x glutamax and 1x penicillin-streptomycin. Lentiviruses were produces by co-transfecting packaging vectors (psPAX2 and pMD2.G) and transfer vector (pCDH-EF1a-MCS-PuroR) in HEK-293T cells. Cells were transduced with lentiviruses for 2 days and selected using 1.5 μg/μl puromycin for 4 days. Mouse and human EZHIP DNA sequences were used in mouse and human cell lines respectively; however, only human amino acid numbers were used to avoid confusion. S2 cells in this study were cultured in Schneider’s Media (Thermo Fisher) containing 10% FBS (Omega Scientific), and 1% antibiotic/antimycotic (Thermo Fisher). Human EZHIP and H3.3 genes was cloned into the pMT-puro vector for copper-inducible expression with the metallothionein (MT) promoter. Transfections were performed with 2 μg plasmid DNA, using Effectene Transfection Reagent for random genome integration, and selected using 2 μg/ml puromycin for approximately three weeks.

### Fly Stocks

All stocks were grown on molasses food at 22°C (room temperature). N-terminally FLAG-tagged EZHIP, EZHP(M406E), H3K27M or H3K27R were cloned into pUASt-attB (DGRC#1419) and integrated into ZH-86Fb on the third chromosome using PhiC31 integrase-mediated recombination into BDSC#24749 with fluorescence marker removed (Best Gene). *en-Gal4, UAS-RFP/CyO* (II) (BDSC#30557) was used to drive expression in the larval wing disc.

### Immunohistochemistry

Crawling third instar larvae were harvested and dissected in pre-chilled (4°C) 1X PBS. Wing imaginal discs were removed from larvae and placed into 1X PBS on ice and fixed in 4% formaldehyde for 30 minutes. Fixed wing imaginal discs were washed in 1X PBS + 0.1% Triton X-100, (PBST) and blocked in PBST + 1% BSA (PAT). After removal of PAT, wing discs were resuspended in PAT + anti-H3K27me3 (1:1600) (Cell Signaling Technology #9733S) and incubated overnight at 4°C, washed in PBST, incubated in PBST + 2% normal goat serum for 10 minutes, followed by incubation with PBST + 2% normal goat serum and goat anti-rabbit DyLight 488 conjugated secondary antibody (1:2000) (Fisher Scientific #35552). Larvae were imaged at 10X using a Nikon Ti2-E epifluorescent microscope.

### Production of stable S2 cell lines for inducible EZHIP expression

S2 cells in this study were cultured at 25°C in Schneider’s Media (Thermo Fisher) containing 10% FBS (Omega Scientific), and 1% antibiotic/antimycotic (Thermo Fisher). FLAG-tagged human EZHIP was cloned into the pMT-puro vector (Addgene #17923). Transfections were performed with 2 μg plasmid DNA, using Effectene Transfection Reagent (Qiagen), and cells were selected using 2 μg/ml puromycin for approximately three weeks. For induction, 5 μM or 10 μM copper sulfate was added to cells at one million cells/ ml density. Cells were incubated for 72 hours and harvested for immunoblot or ChIP.

### FLAG affinity Purification

~80 million cells were homogenized in hypotonic lysis buffer (15 mM HEPES pH 7.9, 4 mM MgCl2, 10 mM KCl, 1 mM EDTA, 8 mM PMSF) to isolate nuclei. Nuclei were resuspended in Buffer-M (15 mM HEPES pH 7.9, 1 mM CaCl2, 30 mM KCl, 1X protease inhibitor cocktail, 8 mM PMSF, 1 mM beta-mercaptoethanol) and treated with 750 units of MNase for 20 min at 37 °C. MNase digestion was quenched and nuclear extract was prepared by adding 10 mM EDTA, 5 mM EGTA, 270 mM KCl, 0.05% Triton X-100). Nuclear extract was incubated with 75 μl of M2 anti-FLAG affinity gel (Sigma A2220) for 2 hours. Beads were washed 5-times with wash buffer (15 mM HEPES pH 7.9, 500 mM KCl, 1 mM EDTA, 0.05% Triton X-100, 8 mM PMSF) and captured proteins were eluted using 300 μg/ml of 3x FLAG peptides. For FLAG affinity purification in the presence of A-395, 1 μM A-395 (or DMSO control) was added to cultured cells for 6 hours before cells were harvested and nuclear extract was prepared. 1 μM A-395 (or DMSO) was added to all buffers throughout the protocol.

### Immunoprecipitation of pre-deposition complexes

Lysate from 40 × 10^6^ HEK-293T cells transduced with H3.3-FLAG-HA transgenes was prepared by resuspension in 3.0 ml lysis buffer [20 mM HEPES pH 7.9, 200 mM KCl, 0.5 mM EDTA, 2 mM MgCl2, 0.2 % Triton X-100, 2× Protease Inhibitor Cocktail (Roche), 2 mM 2-mercaptoethanol, 1 mM benzamidine, 0.4 mM PMSF, 300 μM S-adenosyl methionine), followed by douncing and separation of the insoluble fraction by centrifugation. Per sample, 30 μl of packed anti-FLAG M2 beads (Sigma) were added to the lysate and incubated rotating at 4 °C for 2 h. Beads were transferred onto microspin columns (Enzymax) and washed three times with wash buffer (20 mM HEPES pH 7.9, 300 mM KCl, 1 mM EDTA, 0.12 % Triton X-100, 0.4 mM PMSF, 1 mM benzamidine, 150 μM S-adenosyl methionine) for 5 min. Finally, samples were eluted with 2 × 25 μl elution buffer [wash buffer supplemented with 500 ng μl^-1^ 3×FLAG peptide (Tufts University Peptide Core Facility)] via incubation for 5 min on ice and centrifugation at 300 g.

### Histone Methyltransferase Assays

200 nM oligonucleosome arrays or 25 μM H3 peptide (18-32) substrates were incubated with 20 nM recombinant PRC2 complex, 4 μM S-adenosyl Methionine (1 μM 3H-SAM; 3 μM cold SAM) and 20 μM H3K27me3 stimulatory peptide in 25 mM Tris pH 8.0, 2 mM MgSO4, 5 mM DTT, 0.4 mM PMSF for 90 min. Reaction was spotted on phosphocellulose membrane (Whatman p81) and dried for 10 min. Filters were washed 3-times with 100 mM NaHCO_3_ for 5 min each, rinsed in acetone and dried for 10 min. Scintillation counting was performed using Tri-Carb 2910 TR liquid Scintillation analyzer (Perkin Elmer). For fluorography, reaction was resolved on 15% SDS-PAGE gel, stained with Coomassie, incubated in Amersham Amplify Fluorography reagent (GE healthcare) for 10 min and dried under vacuum. Films capturing fluorography signal were developed after 24-48 hours. Experiment specific details are in figure legend.

### Chromatin Immunoprecipitation

~20 million mammalian cells or ~80 million S2 cells were cross-linked with 0.8% paraformaldehyde for 8 min at room temperature and quenched with 0.2 M glycine. Cells were lysed by resuspending in lysis buffer (50 mM HEPES pH 7.9, 140 mL NaCl, 1 mM EDTA, 10% glycerol, 0.5% NP40, 0.25% Triton X-100, 0.8 mM PMSF). Nuclei were washed once and resuspended in digestion buffer (50 mM HEPES pH 7.9, 1 mM CaCl2, 20 mM NaCl, 1x protease inhibitor cocktail, and 0.8 mM PMSF), and treated with 200 units of MNase for 10 min. Reaction was quenched by adding 10 mM EDTA, 5 mM EGTA, 80 mM NaCl, 0.1% sodium deoxycholate, 0.5% N-lauroyl sarcosine. Mono-nucleosomes were solubilized by sonication using covaries S220 (160 peak incidental power, 5% duty factor, 200 cycles/burst, 45” ON-30” OFF) 3-times. 1% Triton X-100 was added to the chromatin and insoluble chromatin was removed using centrifugation. Chromatin was dialyzed against RIPA buffer (10 mM Tris pH 7.6, 1 mM EDTA, 0.1% SDS, 0.1% sodium deoxycholate, 1% Triton X-100) for 2 hours. Chromatin concentration was measured using qubit and spike-in chromatin was added at 1:40 ratio. Chromatin was incubated with primary antibodies overnight. Antibodies was captured using Dynabeads for 4 hours and washed 3x using RIPA buffer, 2x using RIPA-NaCl and 2x with LiCl buffer. Chromatin was eluted in 10 mM Tris, 1 mM EDTA, and 1% SDS, incubated with proteinaseK, and RNaseA and DNA was purified using PCR purification columns. For reChIP, chromatin was eluted using a competing 3x-FLAG peptide for 30 min at 10°C. Eluate was diluted 10-times with RIPA buffer and incubated with EZH2 antibody overnight. Final washes and elution were carried out as described for conventional one-step ChIPs. Eluted DNA was diluted 1:50 for qPCR analysis. Sequencing libraries were prepared using NEB Next Ultra kit. ChIPs were performed in at least two independent replicates with similar results, at least one replicate was sequenced using NGS, both replicates were used for qPCR but data in the figures only represent technical replicates; ChIP-qPCR results are displayed as barchart (mean ± SE), p-values were determined by paired, non-parametric t-test.

### ChIP-Sequencing analysis

Reads that passed quality score were aligned to mouse (mm9) or human (hg19) or drosophila (dm6) genomes using bowtie 1 with default parameters. Sample normalization factor was determined as Rx = 10^6 / (total reads aligned to exogenous reference genome). Sam files were converted to bam files using samtools. Bigwig files were generated using deeptools and Rx scale-factor was used for sample normalization. Residual H3K27me3 sites in cells expressing EZHIP or H3 K27M were determined as peaks found in two independent ChIP-Seq experiments. Spreading sites were determined by subtracting EZH2 peaks from broad H3K27me3 peaks present in control samples (EZHIP M406E or H3 K27R). For EZH2 reChIP-Seq analyses, aligned paired-end sequencing reads were filtered by fragment length < 400 bp. Samples were normalized by RPKM and corresponding FLAG ChIP (H3 K27M ChIP). EZH2 reChIP in MEFs not expressing H3 K27M transgene was used as negative control. No conclusion about the amplitude of reChIP signals were drawn in the manuscript. Deeptools was used for data visualization. Peaks were called using mosaics-HMM (typically using FDR= 0.01, maxgap= 2-10K, minsize= 1K). Statistical analysis was performed using R.

For identification of potential PRC2 binding sites in S2 cells, regions containing disproportionate retention of H3K27me3 were determined as the difference in H3K27me3 RPKM enrichment in control and cells expressing EZHIP. Bins with change in H3K27me3 < 10 within 5KB were merged and regions with delH3K27me3 < 500 were removed. Finally, ATAC-Seq peaks within these regions with residual H3K27me3 were defined as potential PRC2 recruitment sites. Annotations of PREs in dm6 genome were obtained from a recent report ^45^.

### ATAC-Sequencing

2 × 10^5^ Drosophila S2 cells were washed once with 1X PBS and then resuspended in 100 μL ATAC lysis buffer (10mM Tris 7.5, 10mM NaCl, 3mM MgCl2, 0.1% NP-40). Cells were centrifuged at 600 × g for 10 minutes at 4°C. The resulting pellet was resuspended in 47.5 μL buffer TD (Illumina 15027866) before adding 2.5 μL Tn5 transposase (Tagment DNA Enzyme, Illumina 15027865) and incubating in 37°C water bath for 30 minutes. The tagmented DNA was immediately purified using MinElute Cleanup Kit (Qiagen 28204) and eluted in 10 μL buffer EB. Tagmented DNA was amplified with 12 cycles of PCR using the NEBNext Hi-Fi 2X PCR Master Mix (NEB M0541) and unique dual index primers. Libraries were purified using a 1.2X ratio of Axygen magnetic beads. 150bp, paire-end sequencing was performed at the University of Wisconsin-Madison Biotechnology Center on the Illumina Nova Seq 6000 platform.

### ATAC-Sequencing analysis

Raw reads were trimmed to remove adapter sequences using NGmerge (Gaspar, 2018, BMC Bioinformatics). Trimmed reads were aligned to the *Drosophila* (dm6) genome using bowtie2 with the following parameters: --very-sensitive, --no-mixed, --no-discordant, −X 5000, −k 2. Only reads with a mapping quality score > 30 that aligned to major chromosomes (2, 3, 4, X, Y) were retained for downstream analysis. In order to enrich for fragments originating from nucleosome-free regions, only fragments < 100 bp were retained. Peak calling was performed on accessible fragments using MACS2 with the following parameters: −f BAMPE --keep-dup all −g 1.2e8 --callsummits.

## Acknowledgments

This research was supported by funding from P01CA196539 (to P.W.L., T.W.M., and N.J.); the Greater Milwaukee Foundation (to P.W.L.), the Sidney Kimmel Foundation (Kimmel Scholar Award to P.W.L.), a startup provided by the Wisconsin Institute for Discovery (to P.W.L.). A.B. is supported by Fonds de Recherche du Québec-Santé. This work was performed within the context of the I-CHANGE consortium and supported by funding from Genome Canada, Genome Quebec, The Institute for Cancer Research of the Canadian Institutes for Health Research (CIHR), McGill University and the Montreal Children’s Hospital Foundation. N.J. is a member of the Penny Cole lab and the recipient of a Chercheur Clinician Senior Award.

## Author Contribution

S.U.J and P.W.L conceptualized the study and wrote the manuscript. S.U.J, A.Q.R., S.D.K., D.H., T.J.D., T.J.G., S.M.L. and E.R.B performed the experiments. S.D., A.S.H. and N.J. helped with NGS samples. N.J., M.M.H, and P.W.L. supervised the experiments. All authors read and edited the manuscript.

---

**Supplementary Figure 1:**
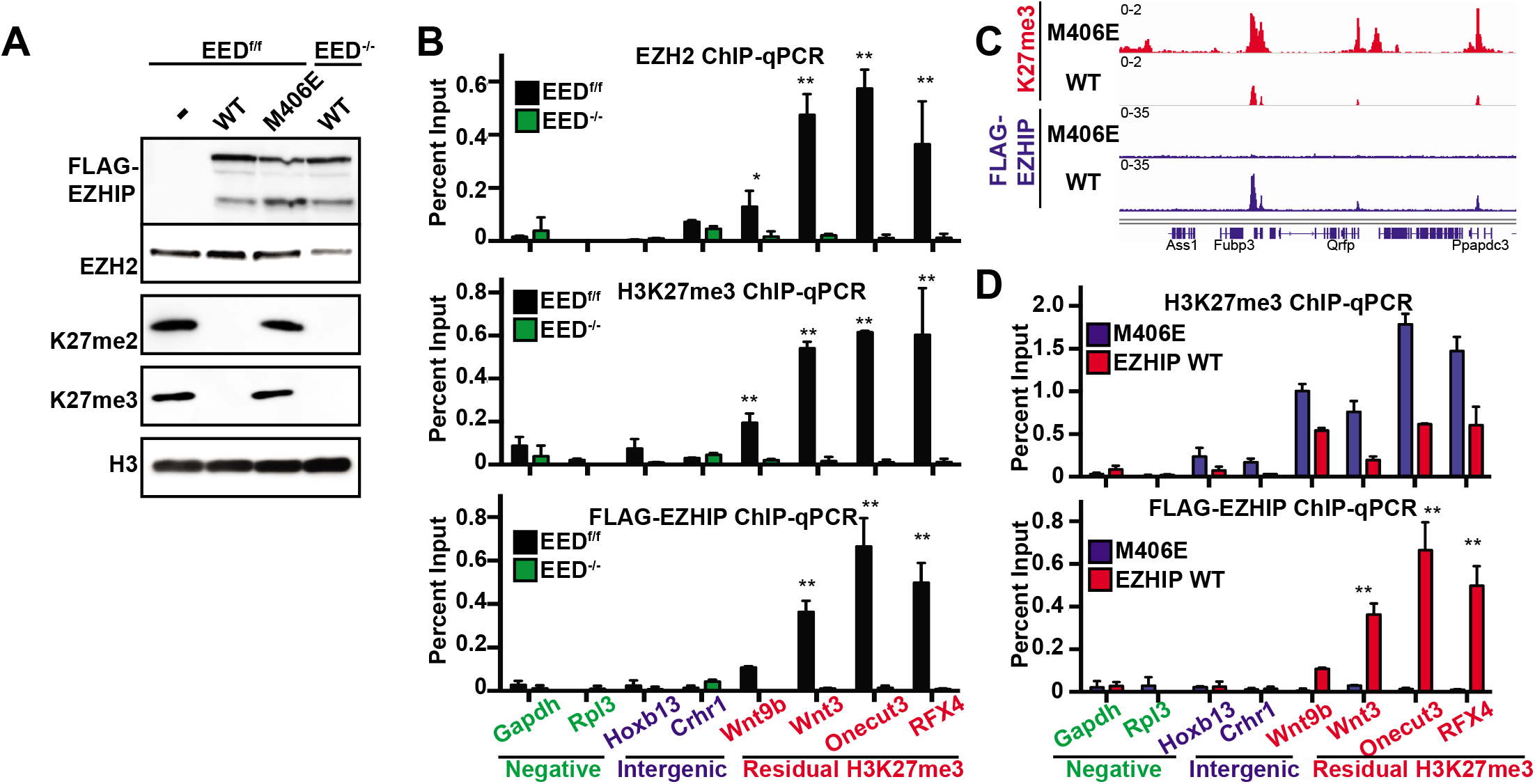
EZHIP colocalizes with PRC2 and residual H3K27me3 *in vivo*: **(A)** Immunoblots of whole cell extracts from cells expressing EZHIP wild-type, EZHIP M406E, and EED knockout cells expressing EZHIP WT. **(B)** ChIP-qPCR of EZH2, H3K27me3 and EZHIP-FLAG ChIPs in cells expressing EZHIP before and after Cre-recombinase mediated EED knockout. **(C)** Genomic profile on H3K27me3 and EZHIP-FLAG ChIPs in cells expressing EZHIP wildtype or M406E. **(D)** ChIP-qPCR analyses of H3K27me3 and EZHIP-FLAG ChIPs in cells expressing EZHIP wildtype or M406E. (**p<0.05)

**Supplementary Figure 2:**
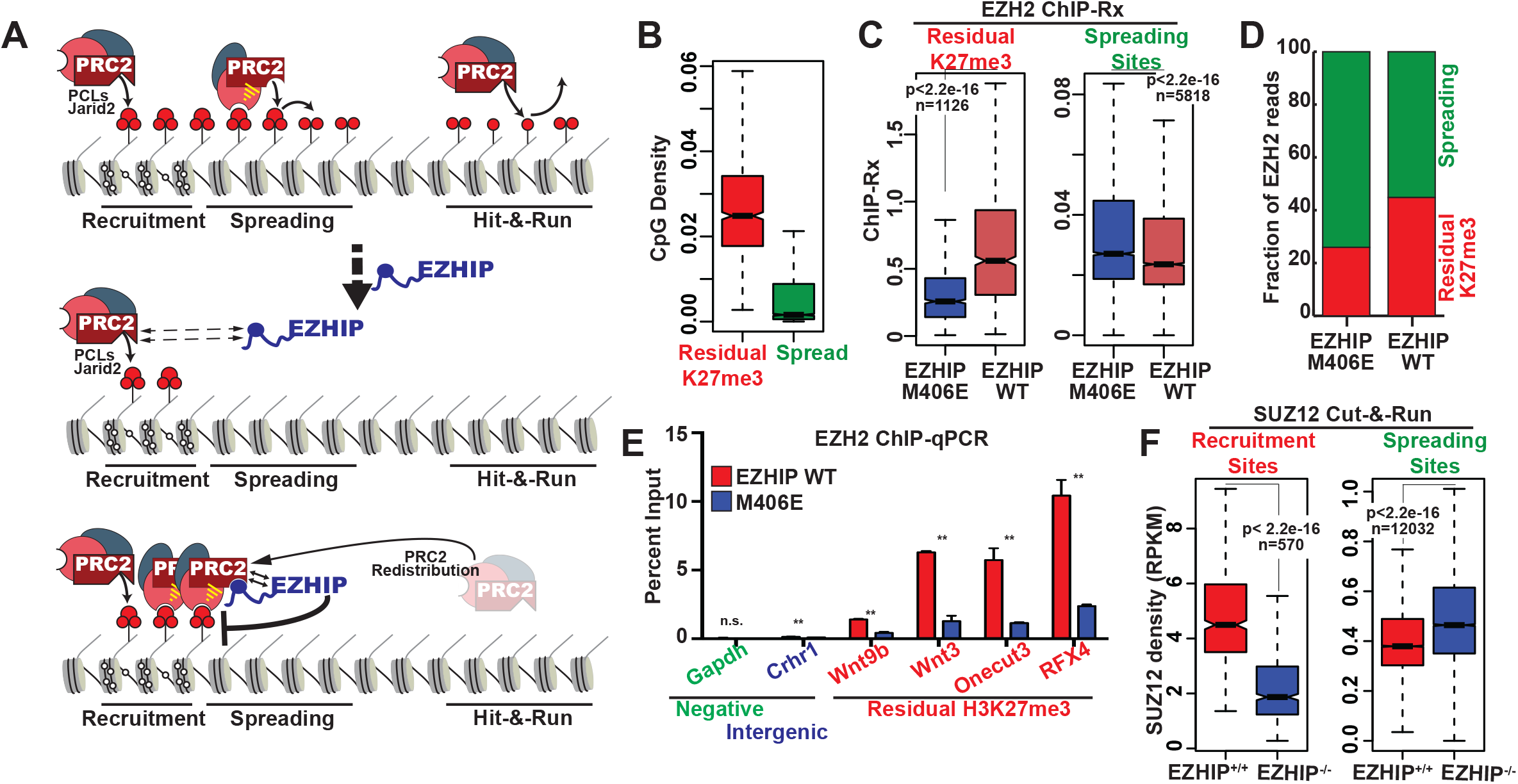
EZHIP sequesters PRC2 at CpG islands containing residual H3K27me3 and prevents PRC2 spreading. **(A)** Schematic depicting the mechanism of EZHIP-mediated PRC2 redistribution. In normal cells, PRC2 is recruited to its high affinity sites at CGIs through polycomb like proteins (PCLs) or Jarid2, where it initiates H3K27me3. Binding of initial H3K27me3 to EED insitigates allosteric-stimulation of EZH2 and PRC2 spreading in cis. Therefore, concentrations of PRC2 at CGIs and spreading sites reaches an equilibrium. In cells expressing EZHIP, PRC2 is able to catalyze initial H3K27me3 since EZHIP has poorer affinity for unstimulated PRC2. However, allosteric stimulation of PRC2 by initial H3K27me3 leads to the formation of a high-affinity EZHIP-PRC2 complex at CGIs. Therefore, PRC2 occupancy is shifted towards CGI away from spreading sites at equilibrium, which results in a disproportionate reduction of intergenic H3K27me3. **(B)** Boxplot displaying CpG densities (total number of CpGs/length of the region) at residual H3K27me3 (red) and spreading (green) sites. **(C)** Boxplot displaying the reference-normalized EZH2 enrichment at recruitment and spreading sites in cells expressing EZHIP WT or M406E mutant. **(D)** Barchart displaying the fraction of EZH2 read densities within the recruitment or spreading sites in cells expressing EZHIP WT or M406E. **(E)** EZH2 ChIP enrichment in cells expressing EZHIP WT or M406E as measured by qPCR (p<0.05**). **(F)** Boxplot displaying the reference-normalized EZH2 enrichment at recruitment and spreading sites in EZHIP^+/+^ and EZHIP^-/-^ U2OS osteosarcoma cells.

**Supplementary Figure 3:**
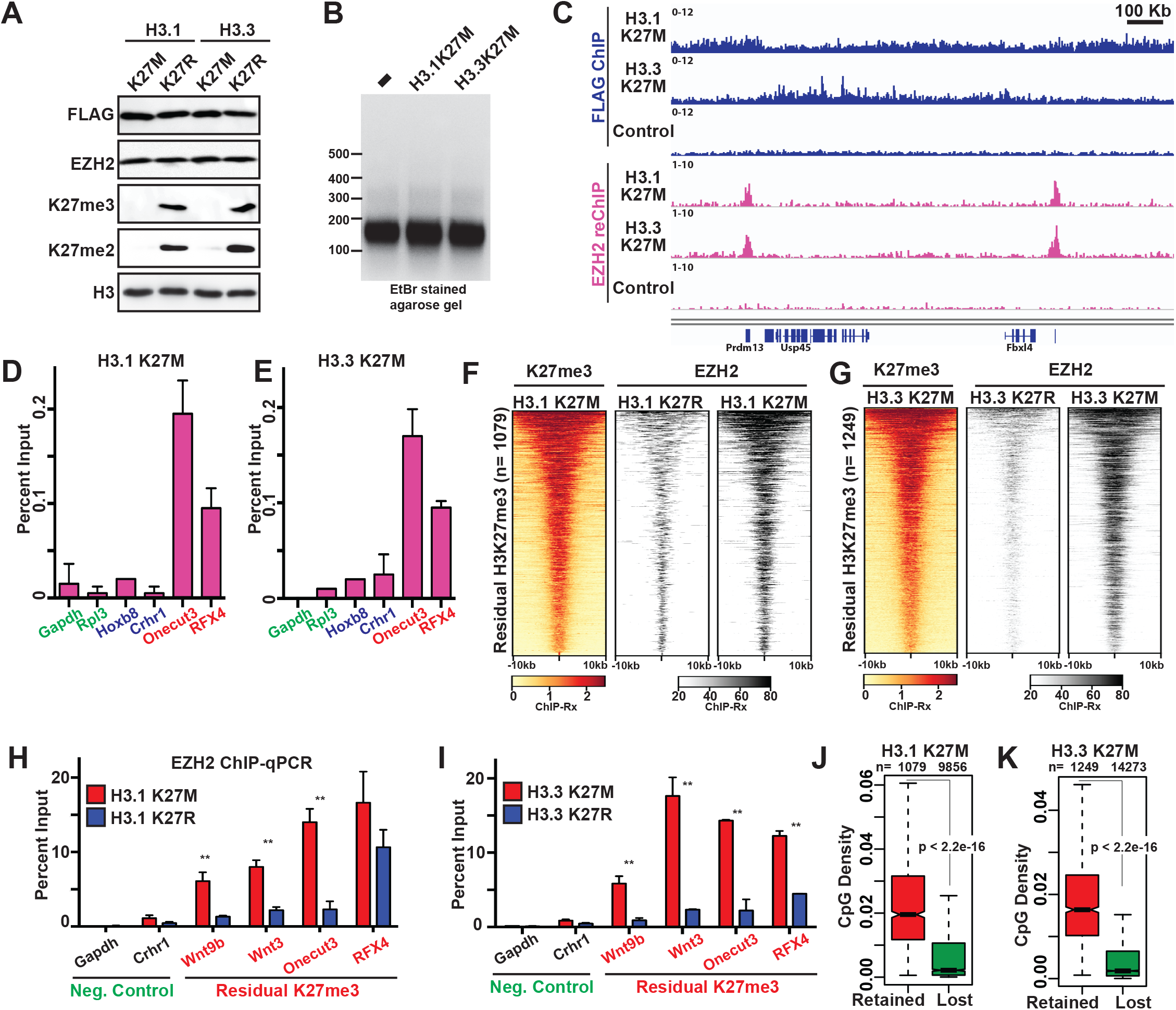
H3K27M sequester PRC2 at CpG islands containing residual H3K27me3. **(A)** Immunoblots from whole cell extracts of MEFs expressing H3.1 or H3.3 K27M and K27R. **(B)** Ethidium bromide stained agarose gel displaying the fragment size distribution of nucleosomes used for FLAG ChIP followed by Ezh2 reChIP experiments. **(C)** Genome browser view of FLAG-EZH2 ChIP-reChIP densities in cells expressing H3.1K27M or H3.3K27M. EZH2 reChIP enrichment was not found in control cells lacking FLAG-tagged H3 transgene. **(D and E)** EZH2 reChIP enrichment at a subset of promoters in cells expressing H3.1K27M **(D)** or H3.3K27M **(E)** as measured by qPCR. **(F and G)** Heatmap displaying EZH2 enrichment at residual H3K27me3 in cells expressing H3.1 K27M and K27R **(F)**, or H3.3 K27M and K27R **(G)**. **(H and I)** EZH2 ChIP enrichment in cells expressing H3.1K27M **(H)** or H3.3 K27M **(I)** at a subset of target loci measured by qPCR (** p<0.05). **(J and K)** Boxplot displaying CpG densities at residual H3K27me3 peaks and regions that displayed loss of H3K27me3.

**Supplementary Figure 4.**
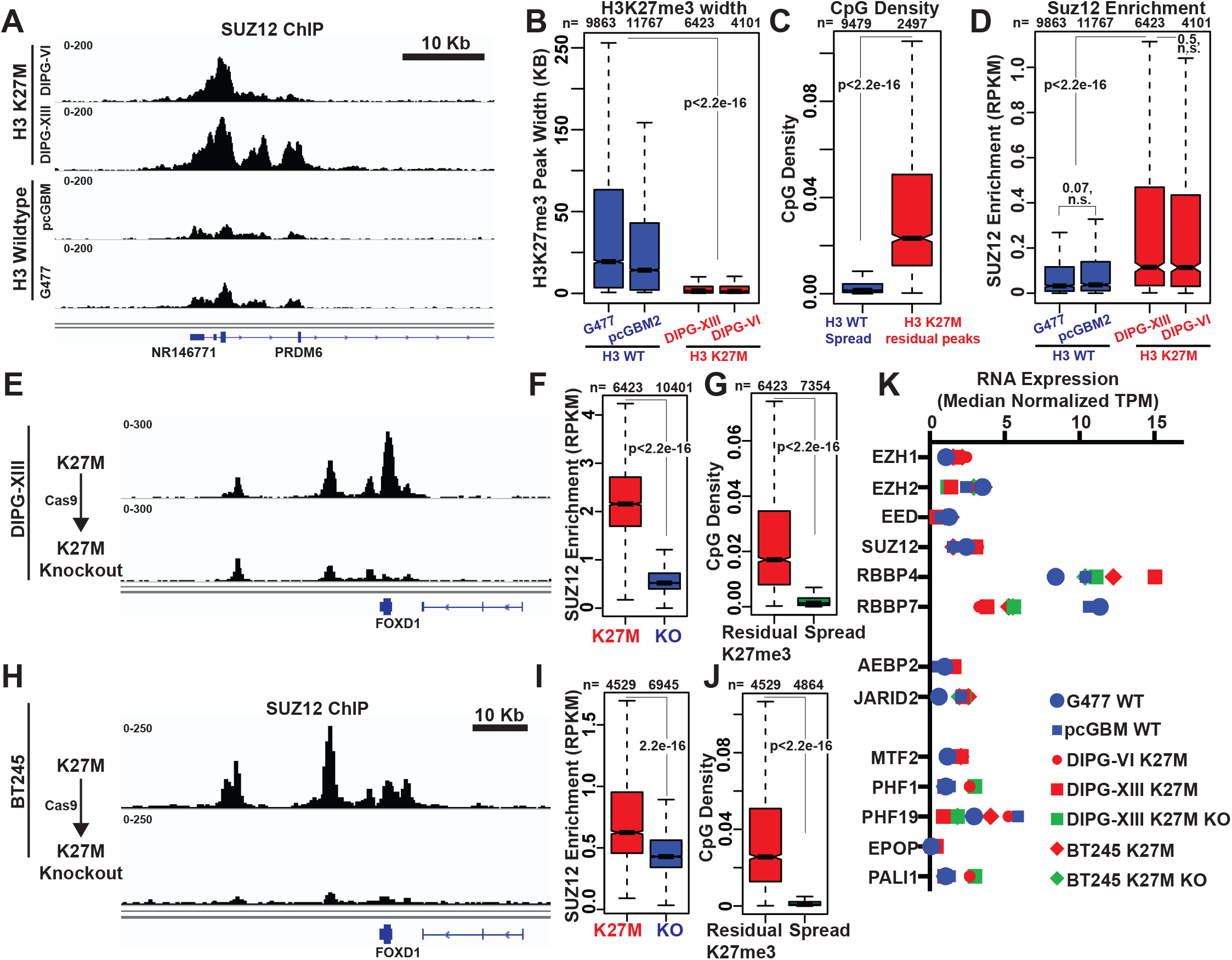
H3 K27M sequesters PRC2 at CpG islands containing residual H3K27me3 in Diffuse Midline Gliomas. **(A)** Genome browser view of SUZ12 occupancies in DMG cell lines containing H3 K27M mutations (DIPG-XIII and DIPG-VI) or H3 WT (G477 and pcGMB2). **(B)** Boxplots displaying the H3K27me3 peak width in DMG cell lines as describes in A. **(C)** Boxplot displaying CpG densities within residual H3K27me3 peaks (red) or spreading sites that lost H3K27me3 (blue). **(D)** Boxplots displaying RPKM normalized SUZ12 enrichment at residual H3K27me3 peaks. **(E and H)** Genome browser view of SUZ12 occupancies in DIPG-XIII **(E)** or BT245 **(H)** cell line containing H3.3 K27M mutation or after cas9-mediated genetic depletion of H3.3 K27M. **(F and I)** Boxplot displaying SUZ12 RPKM enrichment at residual H3K27me3 peaks in DIPG-XIII cells **(F)** or BT245 cells **(I)** containing H3.3 K27M or H3.3 K27M Knockout cells. **(G and J)** Boxplot displaying CpG densities wihin residual H3K27me3 peaks or at spreading sites that gained H3K27me3 upon H3.3 K27M knockout in DIPG-XIII **(G)** or BT245 cells **(J)**. **(K)** Plot displaying the RNA expression of PRC2 subunits in DMGs containing H3 WT (blue) or H2 K27M mutations (red) or H3.3 K27M knockout cells (green). None of the PRC2 subunits were differentially expressed in H3 K27M mutant tumors or upon H3 K27M knockout.

**Supplementary Figure 5.**
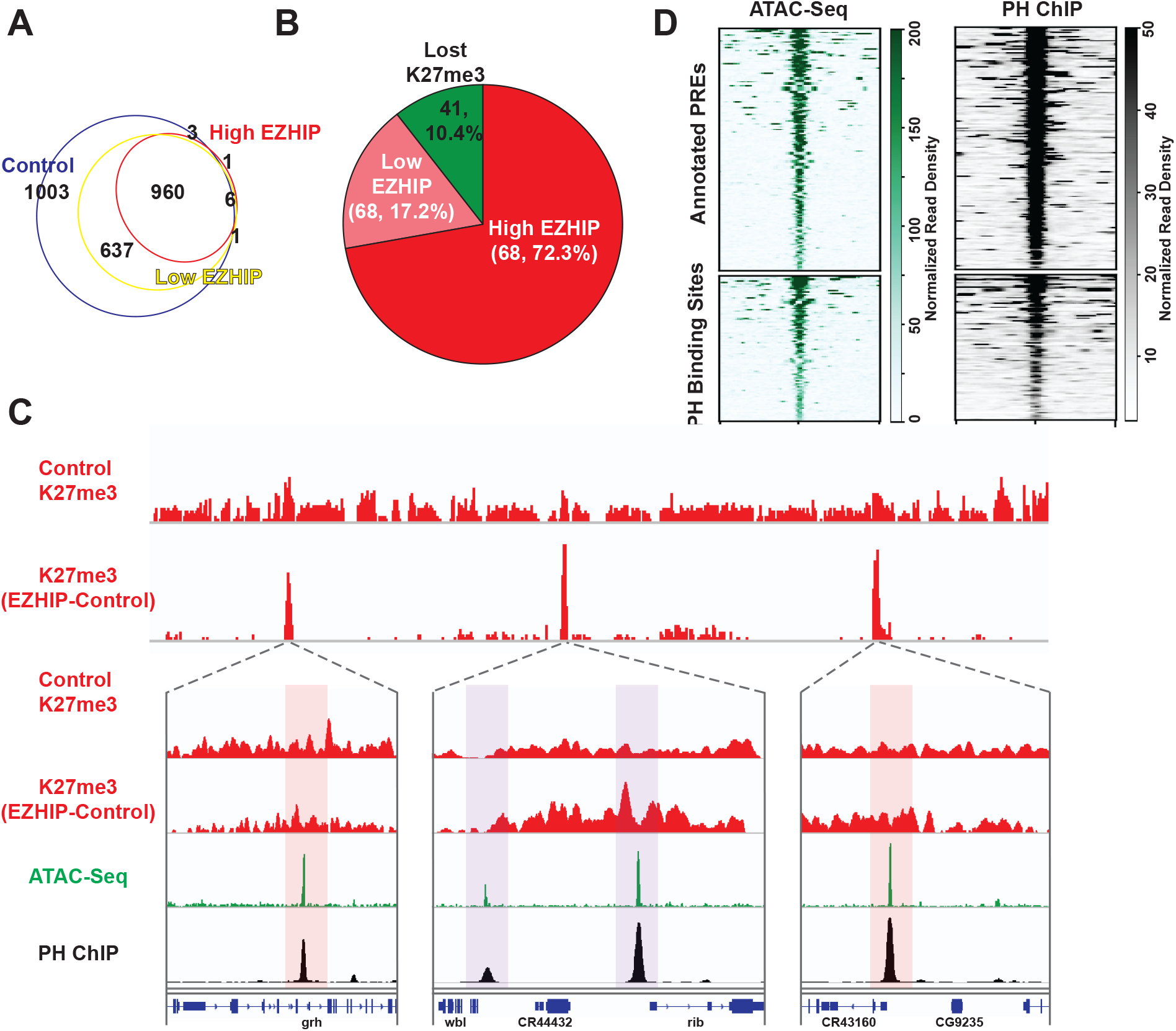
Human EZHIP reduces global H3K27me3 but retains residual H3K27me3 at Polycomb-recruitment sites in Drosophila S2 cells. **(A)** Venn diagram displaying the overlap between H3K27me3 peaks found in control S2 cells (blue) or cells induced to express EZHIP at 5 μM (yellow) and 10 μM (red) concentrations. **(B)** Density of annotated PREs within the residual peaks of H3K27me3 in S2 cells expressing EZHIP at 10 μM Cu^2+^. **(C)** Genome browser view of H3K27me3 ChIP, H3K27me3 (EZHIP-Control), ATAC-Seq and PH ChIP-Seq profiles (related to figure **4G**). Previously annotated PREs and other polycomb recruitment sites are highlighted in red and purple respectively. **(D)** Heatmap displaying ATAC-Seq and PH ChIP-Seq read enrichments at polycomb recruitment sites identified in Figure 4D.

